# Prostaglandin E_2_ induction by cytosolic *Listeria monocytogenes* in phagocytes is necessary for optimal T-cell priming

**DOI:** 10.1101/2021.03.23.436585

**Authors:** Courtney E. McDougal, Zachary T. Morrow, Seonyoung Kim, Drake Carter, David M. Stevenson, Daniel Amador-Noguez, Mark J. Miller, John-Demian Sauer

**Author notes:** Corresponding Author (JDS).

## Abstract

*Listeria monocytogenes* is an intracellular bacterium that elicits robust CD8^+^ T-cell responses. Despite the ongoing development of *L. monocytogenes*-based platforms as cancer vaccines, our understanding of how *L. monocytogenes* drives robust CD8^+^ T-cell responses remains incomplete. One overarching hypothesis is that activation of cytosolic innate pathways is critical for immunity, as strains of *L. monocytogenes* that are unable to access the cytosol fail to elicit robust CD8^+^ T-cell responses and in fact inhibit optimal T-cell priming. Counterintuitively, however, activation of known cytosolic pathways, such as the inflammasome and type I IFN, lead to impaired immunity. Here, we describe a cytosol-dependent response that is critical for immunity to *L. monocytogenes*, namely production of prostaglandin E_2_ (PGE_2_) downstream of cyclooxygenase-2 (COX-2). Vacuole-constrained *L. monocytogenes* elicit reduced PGE_2_ production compared to wild-type strains in macrophages and dendritic cells *ex vivo*. *In vivo,* infection with wild-type *L. monocytogenes* leads to 10-fold increases in PGE_2_ production early during infection whereas vacuole-constrained strains fail to induce PGE_2_ over mock-immunized controls. Mice deficient in COX-2 specifically in Lyz2^+^ or CD11c^+^ cells produce less PGE_2_, suggesting these cell subsets contribute to PGE_2_ levels *in vivo,* while depletion of phagocytes with clodronate abolishes PGE_2_ production completely. Taken together, this work identifies the first known cytosol-dependent innate immune response critical for generating CD8^+^ T-cell responses to *L. monocytogenes,* suggesting that one reason cytosolic access is required to prime CD8^+^ T-cell responses may be due to induction of PGE_2_.

**Author summary:** *L. monocytogenes* is an intracellular bacterial pathogen that generates robust cell-mediated immune responses. Due to this robust induction, *L. monocytogenes* is used as both a model to understand how CD8+ T-cells are primed, as well as a platform for cancer immunotherapy vaccines. *L. monocytogenes* must enter the cytosol of an infected host cell to stimulate robust T-cell responses, however, which cytosolic innate pathway(s) contribute to T-cell priming remains unclear. Here, we define COX-2 dependent PGE_2_ production as the first cytosol-dependent innate immune response critical for immunity to *L. monocytogenes*. We found that *ex vivo* PGE_2_ production by macrophages and dendritic cells is partially dependent on cytosolic access, as vacuole-constrained strains of *L. monocytogenes* elicit reduced PGE_2_. *In vivo,* cytosolic access is essential for PGE_2_ production. *L. monocytogenes* elicits a 10-fold increase in PGE_2_ production, whereas strains of *L. monocytogenes* that cannot access the cytosol fail to elicit PGE_2_ compared to mock immunized mice. Furthermore, CD11c^+^ and Lyz2^+^ cells contribute to PGE_2_ production *in vivo*, as mice deficient in COX-2 in these cell subsets have impaired PGE_2_ production. Taken together, our work identifies the first known cytosol-dependent pathway that is critical for generating immunity to *L. monocytogenes*.

## Introduction

*Listeria monocytogenes* is a Gram-positive, intracellular pathogen that elicits robust CD8^+^ T-cell responses. Due to this robust response, *L. monocytogenes* has been used for decades as a model to understand how CD8^+^ T-cell responses are primed[1]. Understanding these responses has become more pressing recently as *Listeria*-based platforms aiming to drive CD8^+^ T-cell responses are in use as cancer immunotherapies[2]. Initial work showed that critical signals promoting *Listeria*-stimulated T-cell responses are provided acutely, as bacterial clearance with antibiotics as early as 24 hours-post infection has minimal impact on the kinetics of CD8^+^ T-cell responses[3]. This work highlights the role of early signals in informing *Listeria-* stimulated cell mediated adaptive responses.

One early signal impacting T-cell responses is the inflammatory environment induced during infection. The importance of the inflammatory milieu on priming T-cell responses has been solidified by multiple groups using antigen-pulsed, matured dendritic cells in combination with non-antigen expressing *L. monocytogenes* as an inflammatory boost[4, 5]. These studies enable discrimination between antigen presentation and inflammation and demonstrate that wild-type *L. monocytogenes* provides an optimal inflammatory milieu to drive T-cell priming[4, 5].

The inflammatory boost provided through wild-type *L. monocytogenes* infection led to increased T-cell responses, whereas use of strains that specifically alter the inflammatory milieu led to suboptimal responses[4, 5]. *L. monocytogenes* activates a number of innate pathways that contribute to the inflammatory milieu. In particular, multiple groups have focused on the role of various cytosolic innate immune pathways, as previous research demonstrated the necessity of cytosolic access in priming cell-mediated immunity[6–8]. *L. monocytogenes* utilizes a cytolysin, listeriolysin O (LLO), to escape from phagosomes directly into the cytosol and LLO-deficient mutants that are unable to access the cytosol inhibit T-cell priming and generate tolerizing immune responses[9, 10]. Despite the importance of cytosolic access for priming T-cell responses, multiple cytosol-dependent innate pathways are counterintuitively detrimental to immunity including STING-dependent type I interferon[11, 12] as well as inflammasome activation[4, 13]. We recently identified an alternative innate pathway, production of the eicosanoid prostaglandin E_2_ (PGE_2_), as important for immunity as mice deficient in PGE_2_ have impaired acute and protective T-cell responses to *L. monocytogenes*[14]. Whether PGE_2_ production is dependent on cytosolic access of *L. monocytogenes* remains unknown as is which cells produce PGE_2_ in response to *L. monocytogenes* infection.

Eicosanoids are lipid mediators of inflammation that have potent biological functions. A major subset of these lipids, including PGE_2_, are derived from arachidonic acid[15]. During inflammation, arachidonic acid is liberated from the membrane by the cytosolic phospholipase A2 (cPLA2) and then further metabolized by a number of enzymes including the P450 epoxygenase, lipoxygenases, and cyclooxygenases (COXs)[15]. During infection, PGE_2_ is produced downstream of COX enzymes, particularly downstream of cyclooxygenase-2 (COX-2)[16]. COX-2 is induced during inflammation and functions to reduce arachidonic acid to prostaglandin H_2_ (PGH_2_)[17, 18]. PGH_2_ is further metabolized into different prostaglandins by terminal prostaglandin synthases. Coupling of COX enzymes with prostaglandin synthases ultimately dictates which prostaglandin will be produced[17, 18]. PGE_2_ specifically is produced by three different terminal synthases, the cytosolic prostaglandin E synthase (cPGES) and microsomal prostaglandin E synthases-1 and -2 (mPGES-1 and mPGES-2)[16, 19]. Of these synthases, mPGES-1 is inducible and associated with infection due to its role in inflammatory responses[16, 19]. For example, mice deficient in mPGES-1 have reduced febrile and pain responses[16, 19]. Previously, we showed that *L. monocytogenes* infection of mice deficient in mPGES-1 or use of a COX-2-specific inhibitor leads to impaired T-cell responses that could be rescued by exogenous dosing of PGE_2_[14]. Together, these data suggest that production of PGE_2_ downstream of COX-2 and mPGES-1 is critical for immunity.

During *L. monocytogenes* infection, the cell types responsible for producing PGE_2_ remain unclear. *L. monocytogenes* is initially captured by a wide range of phagocytic antigen presenting cells (APCs) in the marginal zone of the spleen[20]. Initially, *L. monocytogenes* highly infects multiple macrophage subsets, including MOMA^+^ metallophilic and MARCO^+^ marginal zone macrophages[20]. Later, *L. monocytogenes* infection transitions to splenic CD11c^+^ and CD11b^+^ cells in the white pulp[20]. Importantly, PGE_2_ is produced at high amounts early in the immune response, starting at four hours post immunization and peaking at twelve hours, early timepoints during which macrophages and dendritic cells are heavily infected[14]. Furthermore, one previous study demonstrated that peritoneal macrophages are capable of producing PGE_2_ after *ex vivo* infection with *L. monocytogenes*[21]. Further analysis is required to elucidate whether macrophages and dendritic cells similarly produce PGE_2_ *in vivo*.

Here, we demonstrated *ex vivo* that macrophages and dendritic cells produce PGE_2_ in response to *L. monocytogenes* infection. Importantly, induction of PGE_2_ *ex vivo* was partially dependent on cytosolic access, as infection of bone marrow-derived macrophages or dendritic cells with vacuole-constrained *L. monocytogenes* led to reduced PGE_2_ compared to wild-type strains. In contrast, *in vivo* PGE_2_ production requires cytosolic access, as infection with LLO-deficient *L. monocytogenes* led to a complete lack of PGE_2_ induction, similar to mock-immunized levels. Lyz2^+^ and CD11c^+^ cells contribute to PGE_2_ production *in vivo,* as deletion of COX-2 selectively in these subsets led to reduced splenic PGE_2_ levels. However, these subsets are not solely responsible for production as a small amount of PGE_2_ remains and this remaining PGE_2_ is sufficient to facilitate optimal T-cell priming. Use of phagocyte-depleting clodronate treatment completely eliminated PGE_2_ production to mock-immunized levels. Taken together, this work identifies, for the first time, a cytosolic-dependent pathway critical for inducing immunity to *L. monocytogenes.* We show that phagocytes, particularly macrophages and dendritic cells, produce PGE_2_ in a cytosol-dependent manner.

## Results

### Unprimed macrophages and dendritic cells upregulate PGE_2_-synthesizing enzymes in response to cytosolic *L. monocytogenes*

We previously demonstrated that immunization of mice with *L. monocytogenes* induces production of PGE_2_, starting at four hours post infection and peaking at twelve hours, and that this transient PGE_2_ production is necessary for optimal T-cell priming[14]. During infection, *L. monocytogenes* infects multiple phagocytic cell populations in the spleen, the majority of which are macrophage and dendritic cell subsets[20]. Initially, *L. monocytogenes* localizes to multiple macrophage subsets[20] and by twelve hours after infection, CD11c^+^ dendritic cells comprise the largest subset of *L. monocytogenes* infected cells[20]. We hypothesized that macrophages and dendritic cells were the subsets producing PGE_2_ due these cells being the predominantly infected cell subsets at these early timepoints post immunization. To determine if *L. monocytogenes* infection induces the genes necessary for PGE_2_ production, we first measured expression of *Pla2g4a* mRNA (encoding cPLA2), a phospholipase that releases arachidonic acid from the cell membrane[15], in bone marrow derived macrophages (BMDMs) and bone marrow derived dendritic cells (BMDCs). BMDMs and BMDCs were infected with *L. monocytogenes* and mRNA was harvested six hours later. We found that cPLA2 expression did not change during *L. monocytogenes* infection (S1 Fig). This result was not surprising, as much of cPLA2 activity is modulated by calcium influx and MAPK phosphorylation rather than transcriptional changes[22]. We next measured mRNA expression of *Ptgs2* (encoding COX-2) and *Ptges* (encoding mPGES-1), encoding enzymes involved in the next steps of PGE_2_ synthesis[16, 19]. In both BMDMs and BMDCs, infection with wild-type *L. monocytogenes* led to an increase in *Ptgs2* expression and, to a lower extent, *Ptges*, suggesting that macrophages and dendritic cells could be capable of synthesizing PGE_2_ (Fig 1A-B). Given that PGE_2_ is necessary for optimal T-cell priming and that immunizing mice with a strain of *L. monocytogenes* that cannot access the cytosol leads to reduced T-cell effector function[9, 10], we hypothesized that impaired T-cell responses to vacuole constrained bacteria may be due to reduced expression of PGE_2_-synthesizing enzymes and ultimately decreased production of PGE_2_. To test this hypothesis, we infected BMDMs and BMDCs with a vacuole-constrained strain of *L. monocytogenes* (*Δhly*, a mutant lacking the pore-forming protein LLO) and assessed expression of *Ptges* and *Ptgs2* mRNA. Consistent with this hypothesis, infection with this strain led to reduced *Ptgs2* expression in BMDMs and BMDCs, suggesting that cytosolic access is required for optimal expression of *Ptgs2* (Fig 1A). Interestingly, infection with *Δhly L. monocytogenes* led to similar levels of *Ptges* expression in both BMDMs and BMDCs (Fig 1B). Taken together, these results suggest that cytosolic access increases *Ptgs2* expression in BMDMs and BMDCs, whereas *Ptges* expression is induced independently of cytosolic access. Additionally, as controls we assessed expression of *Ifnb1* (encoding IFN-β) and *Il1b* (encoding IL-1β) in BMDMs and BMDCs. As expected, *Ifnb1* was expressed only during infection with cytosolic, wild-type *L. monocytogenes* in both cell subsets (S1 Fig), where *Il1b* was induced by TLR signaling during infection with both wild-type and *Δhly L. monocytogenes* infection (S1 Fig).

**Fig 1.**
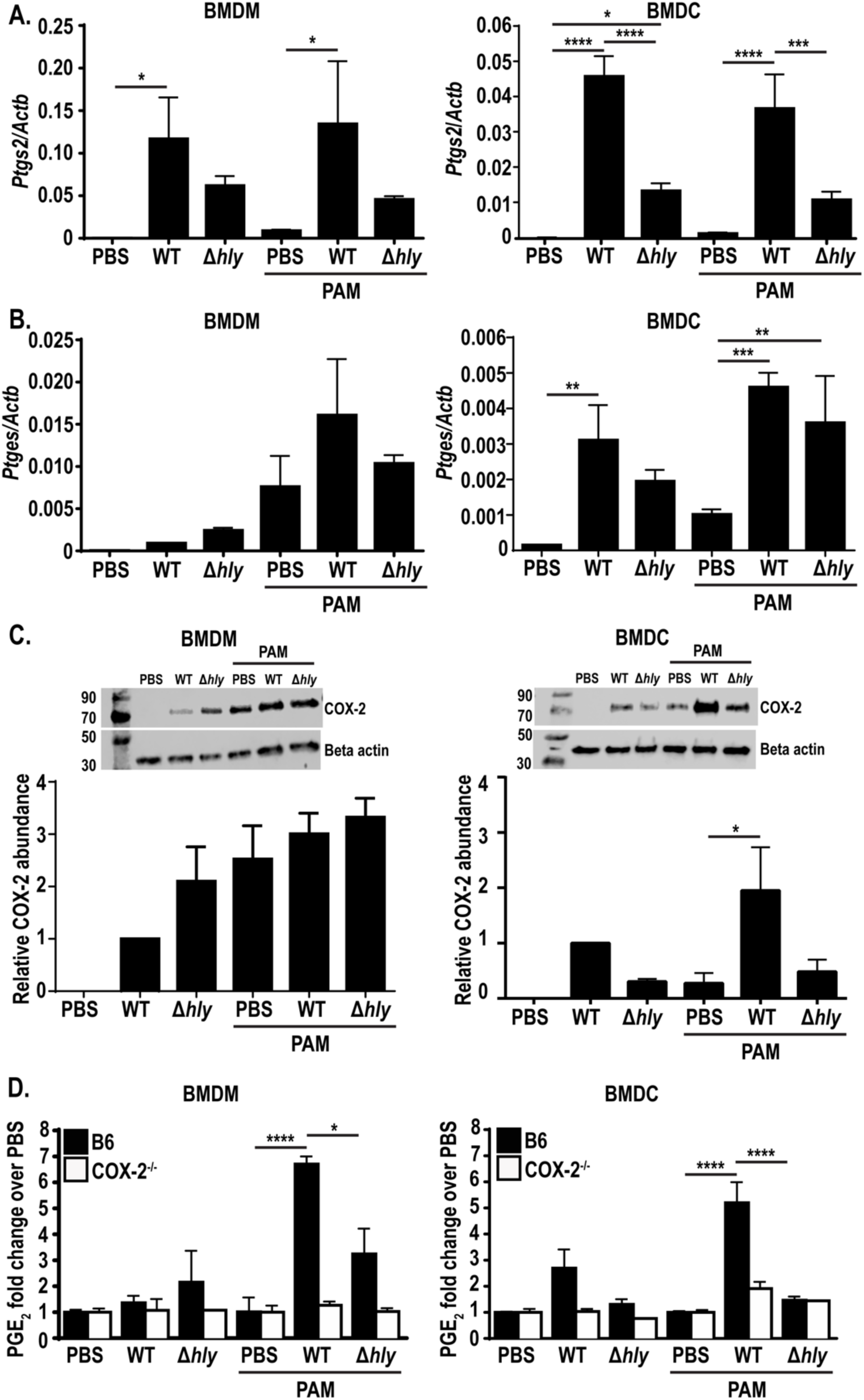
PAM-primed BMDMs and BMDCs express PGE_2_ after cytosolic infection with *L. monocytogenes*. Wild-type or COX-2^-/-^ BMDMs or BMDCs were infected with indicated strains of *L. monocytogenes* at an MOI of 10 *+/-* the TLR2 agonist PAM3CSK4 and assessed 6hpi for expression of *Ptgs2* (encoding COX-2) and *Ptges* (encoding mPGES-1) by qRT PCR (A-B) or COX-2 protein by western blot (C). Supernatant was harvested and assessed for PGE_2_ by mass spectrometry (D). Mass spectrometry data was normalized to d-PGE_2_ and fold change is relative to PBS treated controls. Data are a combination of at least two independent experiments (A,B, D, and western blot quantification), or a representative of at least two independent experiments (western blot image). Significance was determined by a one-way ANOVA with Bonferroni’s correction. \**p* < 0.05, \*\**p* < 0.01, \*\*\**p* < 0.001, \*\*\*\**p* < 0.0001

Given that *Ptgs2* expression was higher during infection with wild-type compared to *Δhly L. monocytogenes*, we next assessed whether the increased transcript in wild-type infection led to increased COX-2 protein expression. To assess the role of cytosolic access on COX-2 protein levels, we infected BMDMs or BMDCs with wild-type or *Δhly L. monocytogenes* and assessed COX-2 protein expression six hours later by western blot. In BMDMs, interestingly, infection with either strain of *L. monocytogenes* led to increased COX-2 protein expression (Fig 1C). In BMDCs, alternatively, *Δhly L. monocytogenes* induced lower levels of COX-2 protein (Fig 1C). This suggests that cytosolic access is required for robust induction of COX-2 protein expression in BMDCs.

Infection of BMDMs and BMDCs with wild-type *L. monocytogenes* led to expression of the genes necessary for PGE_2_ production (Fig 1A-C). To assess whether these cells could utilize these enzymes to produce PGE_2_, we assessed PGE_2_ production in culture supernatant by mass spectrometry. Surprisingly, supernatant from both BMDMs and BMDCs had no detectable PGE_2_ compared to PBS-treated controls, both during infection with wild-type or *Δhly L. monocytogenes* (Fig 1D). This suggests that either enzyme expression was not high enough to induce detectable PGE_2_, or there may be additional post transcriptional modifications required for enzyme activity. Analysis of PGE_2_ from BMDMs or BMDCs deficient in COX-2 had no detectable PGE_2_, as expected (Fig 1D).

### Primed BMDMs and BMDMs produce PGE_2_ during cytosolic *L. monocytogenes* infection

The lack of PGE_2_ produced by BMDMs and BMDCs in response to *L. monocytogenes* infection was surprising given the upregulation of *Ptgs2* and *Ptges* transcript. Other innate pathways, such as the inflammasome, require a priming step in order to induce optimal activation. We hypothesized that macrophages may similarly require additional stimulation in order to produce PGE_2_. To test this hypothesis, we treated BMDMs and BMDCs overnight with the TLR2 agonist PAM3CSK4 (PAM) before infection with wild-type and *Δhly L. monocytogenes* and again analyzed transcript expression. PAM alone induced a small amount of expression of *Ptgs2* expression in both BMDMs and BMDCs (Fig 1A). Infection with wild-type *L. monocytogenes* led to a significant increase in expression that was less robust in *Δhly L. monocytogenes*-infected cells, similar to the effect seen in unprimed cells (Fig 1A). *Ptges* expression, alternatively, had a larger increase in transcript expression during PAM-priming, both during wild-type and *Δhly L. monocytogenes* infection of BMDMs and BMDCs (Fig 1B). Furthermore, PAM treatment alone induced expression of *Ptges* similar to that induced during infection in BMDMs (Fig 1B). Taken together, these data suggest that cytosolic access accentuates expression of *Ptgs2*, where TLR signaling alone is sufficient to induce *Ptges* expression. We also assessed expression of *Pla2g4a* in TLR-primed BMDMs and BMDCs and, similar to unprimed cells, saw no changes in expression (S1 Fig). Additionally, *Ifnb1* transcript was again induced during cytosolic infection, where *Il1b* expression was induced during PAM-treatment alone, as well as during infection with wild-type or *Δhly L. monocytogenes* (S1 Fig).

As *Ptgs2* expression was also dependent on cytosolic access in primed BMDMs and BMDCs, we next assessed expression of COX-2 protein in primed cells infected with wild-type or *Δhly L. monocytogenes* by western blot. Similar to unprimed cells, BMDMs had similar levels of COX-2 protein during infection with wild-type or *Δhly L. monocytogenes* (Fig 1C). In BMDCs, alternatively, COX-2 protein expression was reduced during infection with *Δhly L. monocytogenes* compared to wild-type infection (Fig 1C), again suggesting that COX-2 protein expression in BMDCs is potentiated by cytosolic access.

We hypothesized that priming BMDMs and BMDCs with PAM would stimulate the cells to produce PGE_2_ during infection with wild-type *L. monocytogenes*. To test this hypothesis, we assessed production of PGE_2_ in the supernatant of primed BMDMs and BMDCs by mass spectrometry. BMDMs and BMDCs were treated overnight with PAM before infection with wild-type and *Δhly L. monocytogenes.* Six hours post-infection, cell supernatant was assessed for PGE_2_. In contrast to unprimed BMDM and BMDCs, wild-type infection of primed cells led to a significant increase in PGE_2_ production compared to PBS-treated controls (Fig 1D). Previous data showed *L. monocytogenes*-stimulated PGE_2_ production in peritoneal macrophages[21, 23]. Our data suggest that priming BMDMs prior to infection induces the cells to behave more like tissue resident macrophages in respect to PGE_2_ production. Furthermore, the ability of BMDMs to produce PGE_2_ provides a tool to efficiently study PGE_2_ synthesis in macrophages during infection. Importantly, maximal PGE_2_ production in primed BMDM and BMDCs was dependent on cytosolic access, as infection with *Δhly L. monocytogenes* led to significantly reduced PGE_2_ levels (Fig 1D). PAM-primed COX-2 deficient BMDMs and BMDCs again led to no PGE_2_ production, solidifying the necessity of COX-2 activity in PGE_2_ production (Fig 1D).

Additionally, we also sought to understand whether PGE_2_ specifically was being induced, or if there was a more broad increase eicosanoid production. To test the hypothesis that *L. monocytogenes* induces production of other eicosanoids, we analyzed production of prostaglandin D_2_ (PGD_2_), thromboxane B_2_ (TXB_2_), and leukotriene B_4_ (LTB_4_). However, we saw no changes production of these eicosanoids by wild-type *L. monocytogenes* (S2 Fig). This interesting observation suggests that macrophages and dendritic cells preferentially induce PGE_2_ in response to infection with cytosolic *L. monocytogenes*.

### Cytosolic access is required for PGE_2_ production *in vivo*

Production of PGE_2_ by TLR-primed BMDMs and BMDCs *ex vivo* is potentiated by cytosolic access. To assess whether *L. monocytogenes* induces PGE_2_ in a cytosol-dependent manner *in vivo*, we infected mice intravenously with wild-type and *Δhly L. monocytogenes* and assessed PGE_2_ levels in the spleen twelve hours post-infection, previously defined as the peak PGE_2_ response to infection[14]. Wild-type *L. monocytogenes* led to an eight-fold increase in PGE_2_ (Fig 2A). Infection with *Δhly L. monocytogenes* strikingly showed no increase in PGE_2_ over mock-immunized controls (Fig 2A). To ensure that the reduced PGE_2_ production was not due to differences in bacterial burdens, mice were infected at a dose of wild-type (10^5^ bacteria) and *Δhly L. monocytogenes* (10^7^ bacteria) that led to comparable burdens (Fig 2B). This shows that the absence of PGE_2_ in *Δhly L. monocytogenes-*infected mice is not just due to reduced bacterial burdens. Taken together, these data highlight that cytosolic access is necessary for *in vivo* induction of PGE_2_.

**Fig 2.**
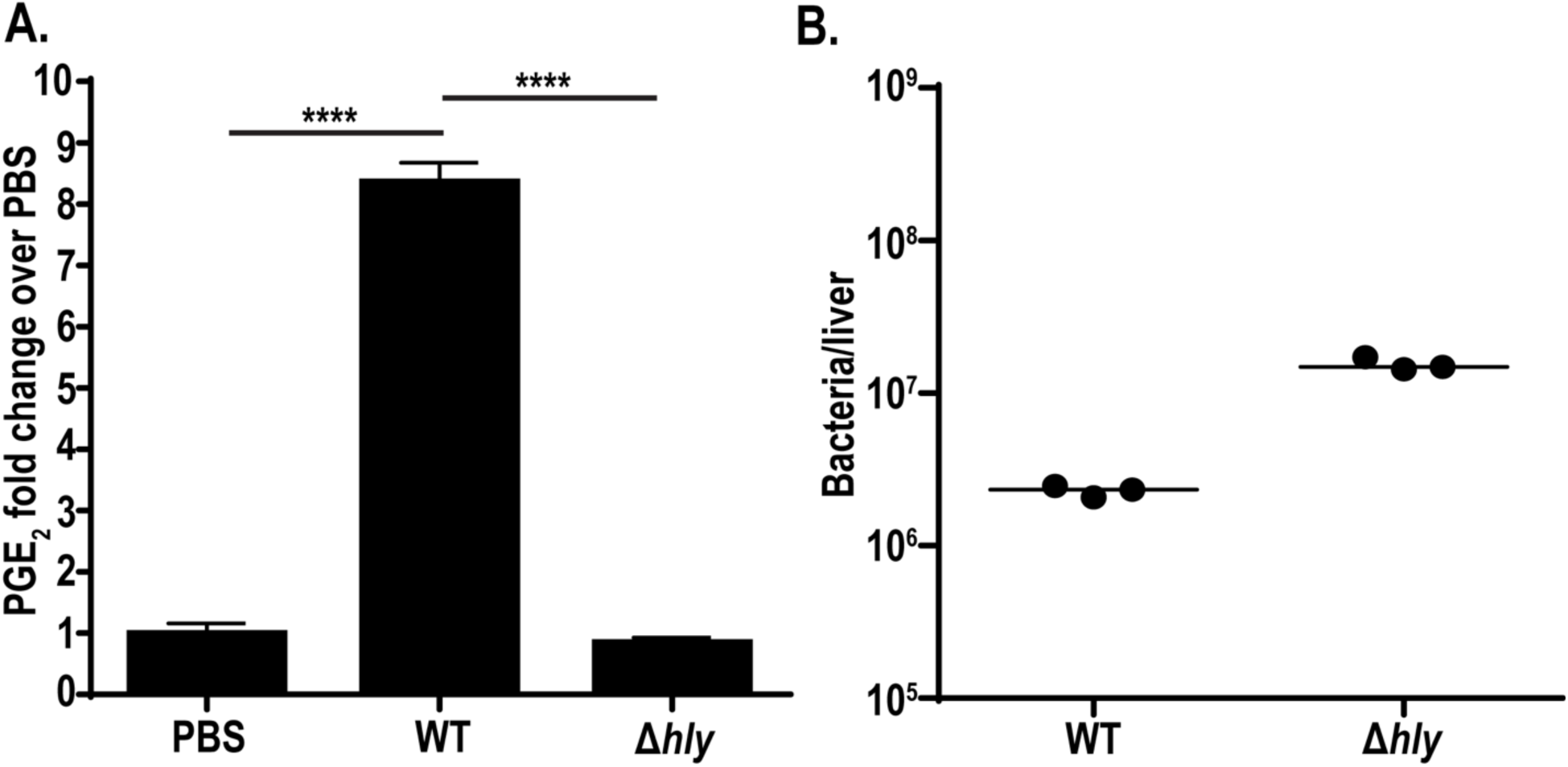
Cytosolic access is necessary for *L. monocytogenes*-stimulated PGE_2_ production *in vivo.* C57BL/6 mice were infected with 10^5^ wild-type or 10^7^ *Δhly L. monocytogenes.* 12hpi spleens were harvested for eicosanoid extraction and mass spectrometry (A) and livers were harvested for bacterial burdens (B). Data are representative of two independent experiments. Mass spectrometry data was normalized to d-PGE_2_ levels and fold change is compared to PBS controls. Significance was determined by a one-way ANOVA with Bonferroni’s correction. \*\*\*\**p* < 0.0001

### CD11c^+^ and Lyz2^+^ cells produce PGE_2_ during *L. monocytogenes* infection *in vivo*

Our data identified PGE_2_ production by macrophages and dendritic cells *ex vivo* (Fig 1D). Furthermore, previous groups have reported that macrophage and dendritic cell subsets are heavily infected early during *in vivo* infection, a timepoint where we have previously detected increases in splenic PGE_2_[14, 20]. From these data, we next hypothesized that macrophages and/or dendritic cells were responsible for producing PGE_2_ *in vivo* that is necessary for optimal T-cell priming. To test this hypothesis, we generated mice deficient in COX-2 selectively in CD11c^+^ cells or Lyz2^+^ cells using the cre/lox system. Mice containing *loxP* sites flanking the COX-2-encoding gene (COX-2^fl/fl^) were crossed with mice expressing the cre recombinase under the CD11c or Lyz2 promoters. COX-2^fl/fl^ CD11c-cre and COX-2^fl/fl^ Lyz2-cre mice (subsequently referred to as CD11c-cre and Lyz2-cre) were immunized with 10^7^ CFU of a live-attenuated, vaccine strain of *L. monocytogenes* (LADD *L. monocytogenes*) currently used in clinical trials as a cancer therapy platform[24]. The LADD strain is deficient in two major virulence genes, *actA* and *inlB*, that retains immunogenicity while making it safe for clinical use[24]. The vaccine strain was used here to enable analysis of T-cell responses in floxed mice as discussed below and induces similar levels of PGE_2_[14]. Immunization of CD11c-cre and Lyz2-cre mice each showed reduced levels of PGE_2_ production, leading to only 60% of the PGE_2_ induced during immunization of control mice (Fig 3A). However, deletion of COX-2 in either CD11c^+^ or Lyz2^+^ cells did not abrogate production to the level of mice globally deficient in mPGES-1 (mPGES-1^-/-^) (Fig 3A). This suggests that CD11c^+^ and Lyz2^+^ cells each contribute to PGE_2_ production and that deletion of COX-2 in either is not sufficient to completely prevent PGE_2_ production. We also assessed PGE_2_ levels in mice deficient in COX-2 selectively in T-cells and observed no reduction of PGE_2_ (S3 Fig). As T-cells are not known to be infected by *L. monocytogenes,* this is consistent with our hypothesis suggesting PGE_2_ production specifically from infected cell subsets.

**Fig 3.**
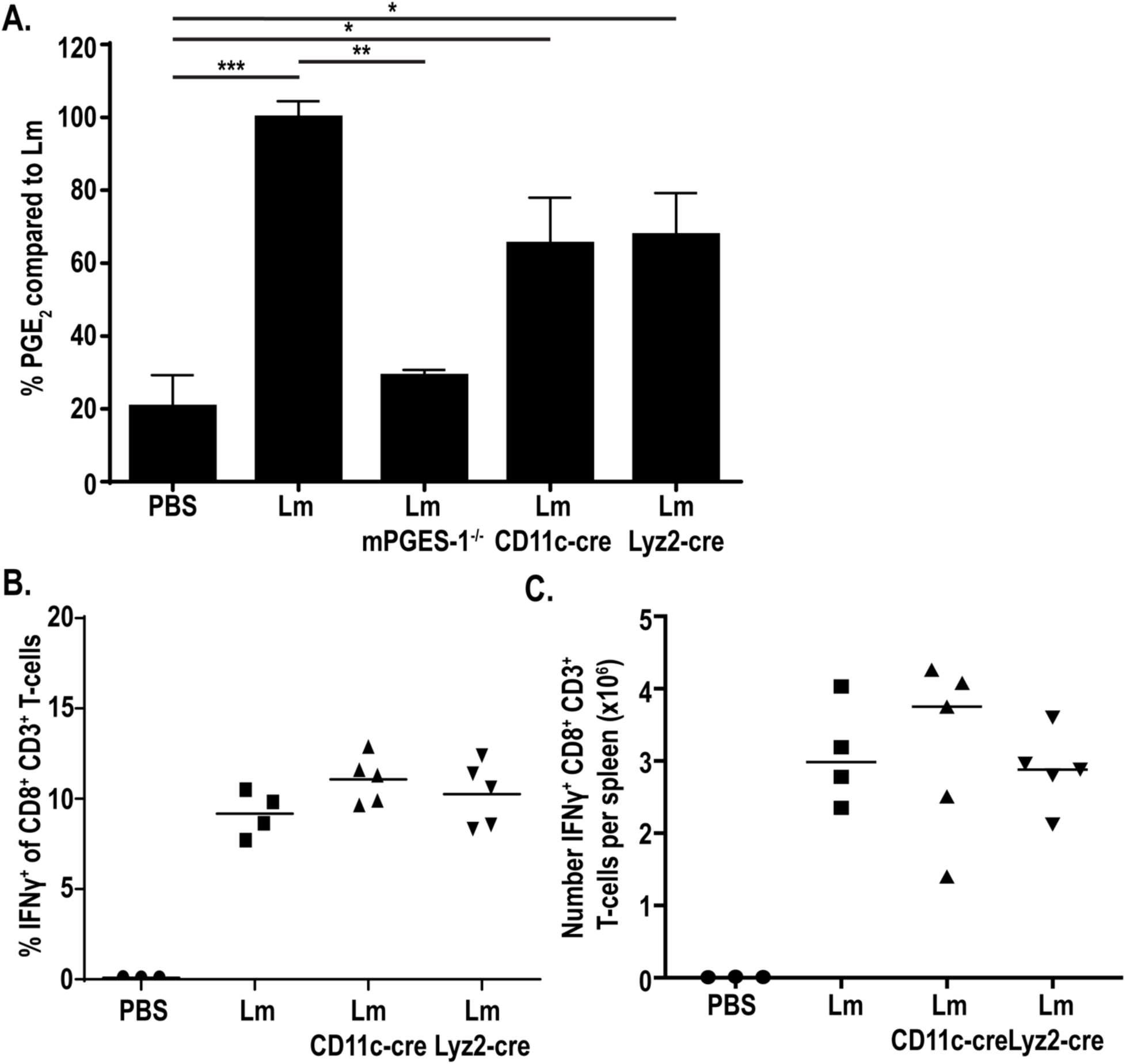
CD11c^+^ and Lyz2^+^ cells contribute to PGE_2_ production *in vivo.* Indicated strains of mice were infected with 10^7^ LADD *L. monocytogenes*. 12hpi spleens were harvested and assessed for PGE_2_ by mass spectrometry. Data was normalized to d-PGE_2_ levels and percent change is compared to *L. monocytogenes*-infected controls (A). Indicated strains of mice were infected with 10^7^ LADD *L. monocytogenes*. 7dpi splenocytes were examined for B8R-specific CD8^+^ T-cell responses. %IFNγ (B) or number IFNγ (C) per spleen was assessed. Data shown are representative of two independent experiments. Significance was determined by a one-way ANOVA with Bonferroni’s correction (A). \**p* < 0.05, \*\**p* < 0.01, \*\*\**p* < 0.001

PGE_2_ is critical for generating optimal T-cell responses in response to *L. monocytogenes,* as immunization of mPGES-1-deficient mice or treatment of mice with a COX-2-specific pharmacological inhibitor leads to impaired T-cell responses[14]. We next hypothesized that the decreased PGE_2_ production in the CD11c-cre or Lyz2-cre mice would be sufficient to similarly impair T-cell responses. To test this hypothesis, we immunized mice with 10^7^ LADD *L. monocytogenes* expressing the model antigens B8R and OVA. Seven days after immunization, splenocytes were isolated, stimulated with B8R or OVA, and production of IFN was assessed by flow cytometry. Despite decreased PGE_2_ production in these mice, T-cell responses were not affected both in percent IFN^+^ as well as number of IFN^+^ T-cells per spleen (Fig 3B-C, S4 Fig) Similarly, the number of antigen-specific T-cells measured by B8R tetramer was unchanged in these mice compared to control mice (S4 Fig). This suggests that the PGE_2_ remaining in these mice was sufficient to prime productive T-cell responses. Due to its short *in vivo* half-life, PGE_2_ asserts its effects locally[25]. It is possible that while global splenic PGE_2_ levels are decreased, the local concentrations of PGE_2_ are sufficient to prime T-cell responses. Taken together, these data suggest that although Lyz2^+^ and CD11c^+^ cells contribute to production of PGE_2_ during *L. monocytogenes* infection, PGE_2_ production by these cells is not necessary, as T-cell responses are not impacted by loss of PGE_2_ production in either subset.

### Deletion of COX-2 in both Lyz2^+^ and CD11c^+^ cells further reduces splenic PGE_2_ levels

Our data showed that single deletions of COX-2 in CD11c^+^ or Lyz2^+^ cells reduced PGE_2_, but not to baseline values. We next hypothesized that PGE_2_ production by either of these subsets individually was sufficient for T-cell priming and that to observe impaired T-cell responses we would have to eliminate PGE_2_ production in both CD11c^+^ and Lyz2^+^ cells. To do this, we crossed the COX-2^fl/fl^ CD11c-cre and COX-2^fl/fl^ Lyz2-cre mice, leading to mice with a COX-2 deletion in both cell subsets (COX-2^fl/fl^ CD11c-cre Lyz2-cre). We assessed the ability of these mice to produce PGE_2_ by mass spectrometry and found that PGE_2_ was further reduced, with about 40% the amount PGE_2_ produced compared to immunized control mice (Fig 4A). This suggests that CD11c^+^ and Lyz2^+^ cells produce the majority of PGE_2_ during immunization with *L. monocytogenes*.

**Fig 4.**
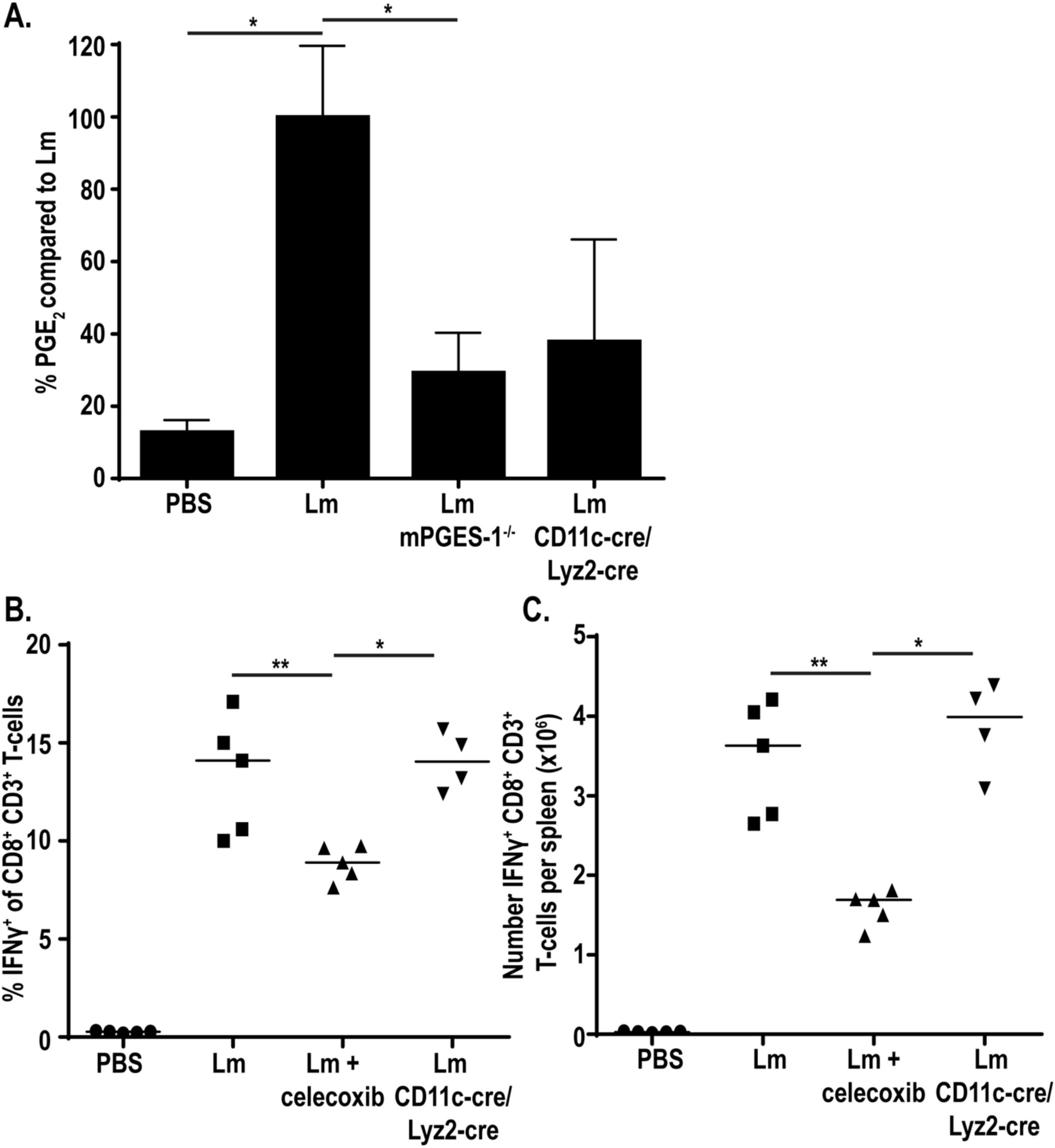
Deletion of COX-2 in both CD11c^+^ and Lyz2^+^ cells further reduces PGE_2_ production. Indicated strains of mice were infected with 10^7^ LADD *L. monocytogenes*. 12hpi spleens were harvested and assessed for PGE_2_ by mass spectrometry. Data was normalized to d-PGE_2_ levels and percent change is compared to *L. monocytogenes*-infected controls (A). Indicated strains of mice were infected with 10^7^ LADD *L. monocytogenes*. 7dpi splenocytes were examined for B8R-specific CD8^+^ T-cell responses. %IFNγ (B) or number IFNγ (C) per spleen was assessed. Data shown are representative of two independent experiments of 3-5 mice per group. Significance was determined by a one-way ANOVA with Bonferroni’s correction (A) or a Mann-Whitney *U* test (B-C). \**p* < 0.05, \*\**p* < 0.01

Due to further reduced PGE_2_ production in our mice deficient in COX-2 in both CD11c^+^ and Lyz2^+^ cells, we next assessed T-cell responses in these mice. Mice again were immunized with 10^7^ vaccine strain of *L. monocytogenes* expressing the model antigens B8R and OVA and assessed for IFN production seven days later. Despite diminished PGE_2_ production, T-cell responses were again not affected, both in percent and number (Fig 4B-C, S5 Fig) Similarly, antigen-specific T-cells measured by B8R tetramer were also unchanged compared to wild-type controls (S5 Fig). This suggest that even the small amount of PGE_2_ produced locally is sufficient to drive T-cell responses.

### Depletion of phagocytes eliminates PGE_2_ production *in vivo*

Our *ex vivo* data highlighted the capability of BMDMs and BMDCs to produce PGE_2_ in response to cytosolic *L. monocytogenes*. However, deletion of COX-2 in Lyz2^+^ and CD11c^+^ cells did not completely abrogate PGE_2_ production *in vivo.* These data led us to hypothesize that other phagocytic cell subsets not effectively targeted by these cre-drivers may be producing the residual PGE_2_, such as marginal zone macrophages (MZMs), metallophilic macrophages, or other CD11b^+^ cells more broadly[26, 27]. To test this hypothesis, we utilized short-term clodronate liposomes to rapidly deplete phagocyte populations in the spleen. Mice were depleted with clodronate liposomes 24 hours prior to immunization with *L. monocytogenes*[28]. Twelve hours post immunization, spleens were harvested and assessed for PGE_2_ by mass spectrometry. Additionally, splenocytes were assessed for CD11b^+^ and CD11c^+^ populations by flow cytometry to confirm clodronate efficacy. Clodronate treatment led to significantly fewer CD11b^+^ cells and a trend for decreased CD11c^+^ cells (S6 Fig). Treatment of mice with clodronate prior to infection with *L. monocytogenes* completely eliminated PGE_2_ production compared to infected control mice (Fig 5A). Importantly, bacterial burdens were equivalent between clodronate and mock-treated mice (Fig 5B). Pretreatment with a control empty liposome, encapsome, actually increased PGE_2_ production compared to infected control mice, potentially due to increased bacterial burdens (Fig 5A-B). Taken together, these data demonstrate that phagocytic cell populations are critical for PGE_2_ production *in vivo* following *L. monocytogenes* immunization.

**Fig 5.**
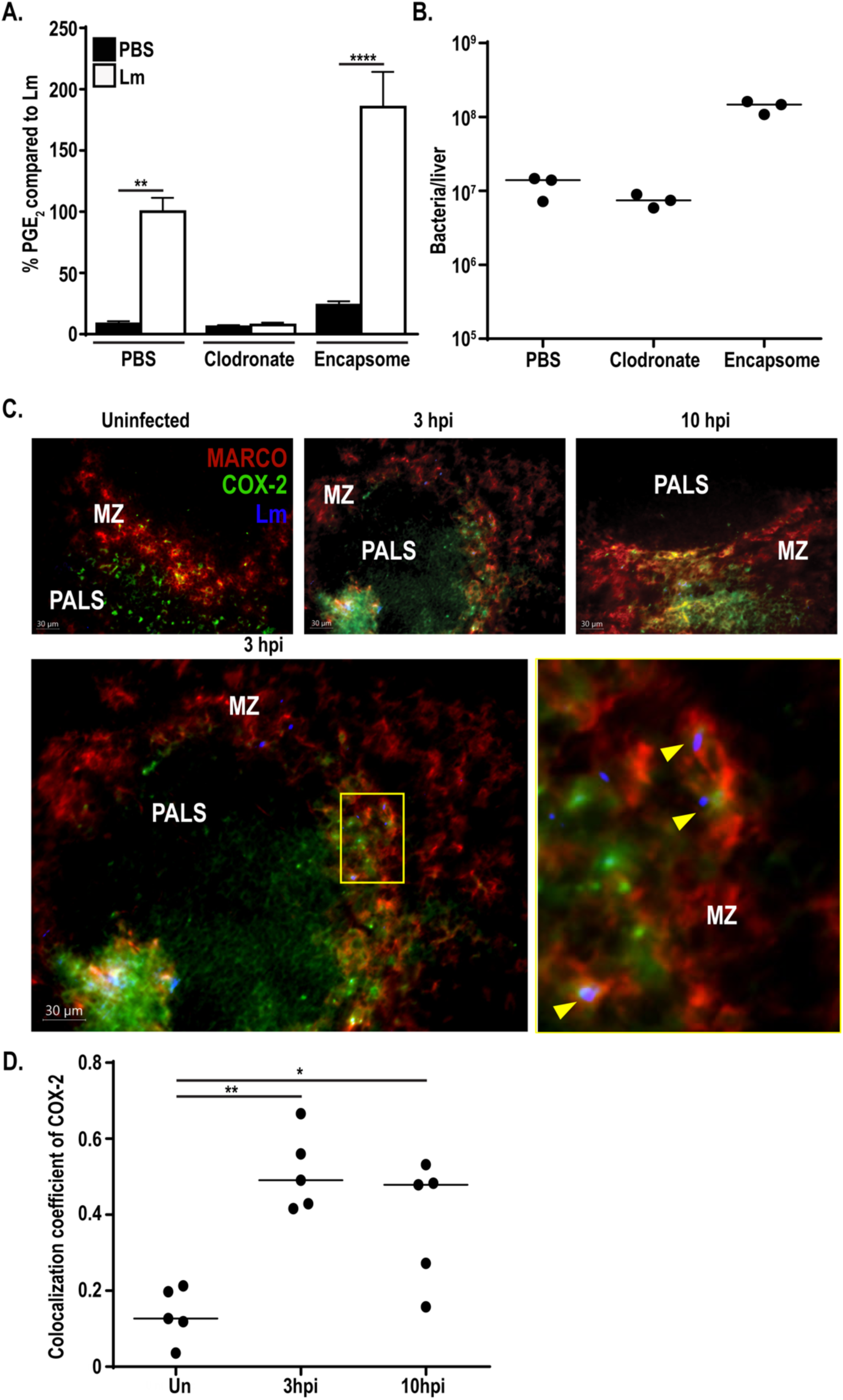
Phagocyte depletion eliminates PGE_2_ production. C57BL/6 mice were dosed with 200µL clodronate, liposome control (encapsome), or PBS 24 hours prior to immunization with 10^7^ LADD *L. monocytogenes*. 12hpi spleens were harvested and assessed for PGE_2_ by mass spectrometry. Data was normalized to d-PGE_2_ levels and percent change is compared to *L. monocytogenes*-infected controls (A). Livers were harvested concurrently and assessed for bacterial burdens (B). C57BL/6 mice were immunized with 10^7^ LADD *L. monocytogenes.* 3 and 10hpi spleens were harvested, cryosections were cut, and sections were stained for *L. monocytogenes* (Lm), COX-2, and MARCO (C). Yellow arrows represent colocalization of *L. monocytogenes*, COX-2, and MARCO (C). Colocalization coefficients (Pearson’s correlation) of COX-2 and MARCO were calculated (D). Data shown are representative of at least two independent experiments. Significance was determined by a one-way ANOVA with Bonferroni’s correction (A) or a Mann-Whitney *U* test (D). \**p* < 0.05, \*\**p* < 0.01, \*\*\*\**p* < 0.0001

Loss of antigen presenting cells through clodronate treatment leads to impaired CD8^+^ T-cell activation, making analysis of T-cell responses in this model not informative[29, 30]. Given this, we alternatively assessed the possibility that other phagocytic cells targeted by clodronate, but not the Lyz2-cre, could contribute to PGE_2_ production. Complete elimination of PGE_2_ production with clodronate treatment suggested that the residual PGE_2_ in the CD11c-cre Lyz2-cre mice was due to a phagocytic cell that was not effectively targeted in these mice. Previous data showed that though the Lyz2-cre used in this study is highly efficient at deletion of *loxP* flanked genes in some macrophage subsets, it is only minimally successful at deleting genes of interest in other subsets, such as MZMs[26]. MZMs, characterized by expression of MARCO, are heavily infected early in *L. monocytogenes* infection[20]. We hypothesized that the residual PGE_2_ we detected in our double CD11c-cre and Lyz2-cre mice may be due to inefficient deletion in macrophage subsets such as these. To assess the role of MZMs in PGE_2_ production, we assessed expression of COX-2 by immunohistochemistry. Mice were immunized with 10^7^ vaccine strain of *L. monocytogenes* and spleens were harvested three and ten hours later. Spleen cryosections were then stained for *L. monocytogenes*, COX-2, and MARCO. Uninfected mice had COX-2 staining in the periarteriolar lymphoid sheath (PALS) with little expression in the marginal zone (MZ) (Fig 5C,-D). As early as three hours post-immunization COX-2 staining was observed in the MZ, with approximately 50% of COX-2 colocalizing with MARCO^+^ cells (Fig 5C-D). Expression of COX-2 in the MZ was maintained at 10hpi, again showing approximately 50% colocalization with MARCO (Fig 5C-D). Furthermore, *L. monocytogenes* colocalized with COX-2 and MARCO expressing cells, suggesting that infected MZMs may be producing PGE_2_ (Fig 5D). Expression of COX-2 suggests that MZMs, or other non-CD11c/Lyz2 expressing phagocytes within the marginal zone, could be capable of producing PGE_2_ *in vivo* and may be contributing to the PGE_2_ remaining in the CD11c-cre Lyz2-cre mice. Taken together, our data suggest that multiple myeloid derived subsets can contribute to PGE_2_ production, including Lyz2^+^ cells, CD11c^+^ cells, and possibly MZMs. Complete reductions in PGE_2_ by depletion of phagocytic cells such as these with clodronate treatment is consistent with our data showing that PGE_2_ is produced from cells infected with cytosolic *L. monocytogenes*.

## Discussion

Cytosolic access is required to effectively generate cell-mediated immunity to *L. monocytogenes*[8–10]. Decades of work has focused on understanding the cytosol-dependent processes necessary for T-cell priming, a topic that has gained interest recently due to use of *L. monocytogenes* as a cancer immunotherapy platform. Our data suggest that one reason cytosolic access is important is to facilitate phagocyte production of PGE_2_, an eicosanoid required to generate optimal CD8^+^ T-cell responses[14]. We showed that PGE_2_ is produced by BMDMs and BMDCs *ex vivo*. Importantly, this pathway requires cytosolic access, as vacuole-constrained *L. monocytogenes* induce lower production of PGE_2_. Furthermore, infection of mice with a vacuole-constrained *L. monocytogenes* strain led to no increase of PGE_2_ over mock immunized controls. Lastly, we showed that Lyz2^+^ and CD11c^+^ cells contribute to PGE_2_ production *in vivo* as deletion of COX-2 in these subsets led to decreased PGE_2_ levels, however other clodronate sensitive phagocyte populations also contribute to PGE_2_ production following *L. monocytogenes* immunization. We demonstrate the first-known innate pathway critical for CD8^+^ T-cell responses that requires cytosolic access by *L. monocytogenes*. This work leads to many new questions including how cytosolic *L. monocytogenes* activates this pathway, how immune cells discriminate which eicosanoid to produce in response to infection, how even small concentrations of PGE_2_ still lead to productive PGE_2_ responses, and how PGE_2_ facilitates optimal T-cell priming.

One intriguing hypothesis is that PGE_2_ synthesis during *L. monocytogenes* infection is driven by an innate cytosolic sensor. *L. monocytogenes* elicits a number of innate pathways that could contribute to differential activation of the PGE_2_-synthesis pathway. One possibility is that induction of type I IFN influences PGE_2_ production. Type I IFN can be induced cytosolically by *L. monocytogenes* through recognition of cyclic diadenosine monophosphate (c-di-AMP). Upon entry into the cytosol, *L. monocytogenes* secretes c-di-AMP through multidrug resistance transporters[31, 32] where it is recognized by either the reductase controlling NF-κB (RECON)[33] or stimulator of IFN genes (STING)[34, 35]. STING activation leads to type I interferon induction[34, 35], and was originally hypothesized to be critical for T-cell responses. Paradoxically, however, type I IFN inhibits cell-mediated immunity to *L. monocytogenes*[12]. Interestingly, there has been well documented crosstalk between the PGE_2_ and type I IFN pathways during infections with other pathogens such as influenza and *M. tuberculosis*[36, 37]. In the context of influenza, Coulombe et al. showed that infection led to upregulation of PGE_2_ and a subsequent decrease in type I IFN[36]. In contrast to *L. monocytogenes,* type I IFN is important in generating cell-mediated immune responses to influenza. Accordingly, diminished type I IFN due to increased PGE_2_ reduces both acute and protective immunity during influenza infection. On the other hand, Mayer-Barber et al. recently showed that inhibition of type I IFN during *M. tuberculosis* infection led to an increased level of PGE_2_ in an IL-1-dependent manner[37]. This correlated with better bacterial control. Due to crosstalk between these two pathways, it seems possible that recognition of c-di-AMP and subsequently upregulation of type I IFN may also be playing a role in PGE_2_ production during *L. monocytogenes* immunization. Analysis of PGE_2_ levels in mice deficient in STING or the type I IFN receptor (IFNAR) could advance understanding of the link between these two pathways. Alternatively, should type I IFN influence PGE_2_ production, use of *L. monocytogenes* strains that have reduced secretion of c-di-AMP and subsequently less type I IFN could be an avenue of further research for immunotherapeutic platforms.

Another cytosolic pathway that may influence PGE_2_ levels during *L. monocytogenes* infection is the inflammasome. Inflammasomes are multiprotein complexes that recognize a wide range of pathogen associated molecular patterns[38–40]. Wild-type *L. monocytogenes* infection leads to a small amount of inflammasome activation, largely through the absent in melanoma 2 (AIM2) inflammasome[41]. The AIM2 inflammasome recognizes cytosolic DNA that is released during bacteriolysis within the cytosol[41–43]. Originally, inflammasomes such as AIM2 were known to have two major downstream effects, the release of proinflammatory cytokines IL1β/IL-18 and the induction of a lytic form of cell death, pyroptosis, characterized by formation of membrane pores by the protein Gasdermin D[44–46]. Seminal work by von Moltke et al. introduced a new downstream effect, the activation of an eicosanoid storm, including PGE_2_[23]. This work, as well as supporting recent work, showed elevated levels of PGE_2_ after inflammasome activation[23, 47]. One possible hypothesis stemming from this work is that induction of membrane pores during pyroptosis leads to calcium influx, activating cPLA2 and releasing arachidonic acid from the membrane. This model would suggest that use of mice deficient in caspase-1 or Gasdermin D would lead to lower levels of PGE_2_ production. Alternatively, infection of *L. monocytogenes* strains that differentially activate inflammasomes would lead to different PGE_2_ production. The role of inflammasomes as well as type I IFN are intriguing avenues to understand signaling pathways driving PGE_2_ production during *L. monocytogenes* infection.

It is also possible that production of PGE_2_ by *L. monocytogenes* is independent of known cytosolic pathways. Identification of other unknown censors could be accomplished by assessing PGE_2_ levels in response to different *L. monocytogenes* mutants. Mutant strains of *L. monocytogenes* that differentially induce PGE_2_ could provide insight as to the cytosolic censors involved. One additional hypothesis is that PGE_2_-production is independent of a cytosolic sensor completely and instead is driven by LLO-mediated pore formation. Though LLO is tightly regulated transcriptionally, translationally, and posttranslationally to be most active in the vacuole, a small amount of LLO may remain active in the cytosol of cells[48–50]. Perhaps, this small amount of LLO induces pore formation in the cell membrane and allows calcium influx, subsequently activating cPLA2. Use of strains that further restrict LLO production in the cytosol, such as new strains that excise *hly* once *L. monocytogenes* has entered the cytosol[51], could help assess the role of LLO-mediated pores on PGE_2_ production.

In addition, our data show an interesting phenotype where BMDCs and PAM-primed BMDMs selectively produce PGE_2_ in response to *L. monocytogenes* infection rather than a global increase in eicosanoid production. Analysis of eicosanoid milieu in cell supernatant show an increase in PGE_2_ production during wild-type *L. monocytogenes* infection, but little to no changes in other eicosanoids such as PGD_2_, TXB_2_, or LTB_4_. This raises the question of how a cell discriminates which eicosanoid is produced in response to different stimuli. The eicosanoid produced in different conditions is dependent on terminal synthases[19]. Therefore, the expression and activity of these synthases determine the resulting eicosanoid milieu. Multiple factors impact expression of different synthases including cytokines, hormones, and microbial products[52]. For example, expression of mPGES-1 can be induced by LPS and prostaglandin D_2_ synthases, though less well understood, can be upregulated by glucocorticoids[53–55]. Use of *L. monocytogenes* strains that differentially activate cytokines or are deficient in different microbial PAMPs could be informative as to which signal specifically leads to enhanced levels of mPGES-1 transcript. In addition, activity of each synthase also may dictate which eicosanoids are produced[52]. Terminal synthase activity can be modulated by posttranslational modification (such as phosphorylation) as well as presence of cofactors (such as ATP and glutathione)[52]. Depletion of essential cofactors during metabolic or oxidative stress could influence the induced inflammatory milieu[52]. The post transcriptional regulation highlights the necessity of assessing endpoint eicosanoid production rather than simply transcript or protein levels, as these other factors influencing activity can alter which eicosanoids ultimately are produced. This is particularly true in our data, as despite seeing upregulation of mPGES-1 transcript during infection of unprimed BMDMs and BMDCs, we failed to see PGE_2_ production. This suggests that perhaps some additional modification is necessary to induce mPGES-1 activity during *L. monocytogenes* infection.

Another pressing question generated from this work is how productive T-cell responses were induced in mice deficient in COX-2 in both CD11c^+^ and Lyz2^+^ cells despite reduced PGE_2_ levels. Here, we show that these mice produce substantially reduced PGE_2_, yet still induce wild-type CD8^+^ T-cell responses. One hypothesis that we explored in this work is that other cell subsets not efficiently targeted by our cre/lox model were still producing PGE_2_. Certain cell subsets such as MZMs do not have effective gene deletion using the Lyz2^+^ promoter to drive cre recombinase expression[26]. For this reason, we hypothesized that subsets such as these may still be producing sufficient levels of PGE_2_ to drive T-cell responses. One way to assess the role of MARCO^+^ MZMs is use of a new cre recombinase-driving promoter, SIGN-R1, developed by Pirgova et al[27]. SIGN-R1 is a lectin binding receptor expressed on MZMs and drives more efficient deletion of genes by the cre/lox system[27]. Generation of triple COX-2 knockout mice that express the SIGN-R1-cre in combination with our reported Lyz2-cre CD11c-cre model could be informative about the role of MZMs in production of PGE_2_. Though we show that Lyz2^+^ and CD11c^+^ cells contribute to PGE_2_, analysis of MZMs and other myeloid cells will further understanding of PGE_2_ production.

The lack of diminished cell-mediated immunity could also be due to local acting effects of PGE_2_. It is possible that even if PGE_2_ levels are below detection at a whole spleen level, certain cells are able to produce PGE_2_ locally in sufficient concentration to drive T-cell responses. More sensitive measures of PGE_2_, such as quantitative mass spectrometry imaging recently developed, would be required to analyze local responses such as these[56]. These novel techniques enable analysis of location of PGE_2_ and other eicosanoids within a spleen and could detect lower concentrations[56]. Similarly, the sensitivity of the receptor PGE_2_ is acting upon during *L. monocytogenes* infection could influence how much PGE_2_ is necessary for inducing a response. PGE_2_ binds primarily to four receptors, EP1-4[57]. EP3 and EP4 are higher affinity receptors (kD ∼1nM compared to 10-15nM for EP1/2)[57, 58]. Should the higher affinity receptors be identified as the important receptors for influencing immunity during *L. monocytogenes* infection, even lower concentrations of PGE_2_ still induced in our Lyz2-cre CD11c-cre model may be sufficient for cell-mediated responses. Further analysis as to relevant receptors and which cells they are expressed on could help elucidate these details.

Lastly, how PGE_2_ facilitates T-cell responses in the context of *L. monocytogenes* immunization remains unknown. In innate immune cells, PGE_2_ influences expression of co-stimulatory and activation markers. PGE_2_ signaling in dendritic cells upregulates the co-stimulatory molecules OX40L and 4-1BBL[59], thereby promoting T-cell proliferation. Similarly, PGE_2_ signaling in macrophages leads to polarization towards a more inflammatory M1 phenotype[60] and aids in activation[61]. Furthermore, PGE_2_ promotes migration of innate cell subsets, leading to enhanced migration towards CCL21[62, 63] and MCP-1[64, 65]. These proinflammatory functions suggest that PGE_2_ may be acting to enhance immunity through its local effects on innate immune cells. PGE_2_ may also be influencing immunity more directly on T-cell subsets, such as through polarization of T-cells towards a Th1 phenotype[66]. Additionally, PGE_2_ leads to higher expression of OX-40L, OX-40, and CD70 directly on T-cells, promoting T-cell interactions and sustaining immune responses[59]. In order to more fully understand how PGE_2_ facilitates T-cell responses to *L. monocytogenes*, a comprehensive analysis of these effects on both T-cells and innate immune cells is required.

We and others have shown that innate immune responses substantially influence cell-mediated immune responses, particularly the inflammatory milieu induced during infection. Here, we present evidence that one pathway critical for immunity, induction of PGE_2_, is dependent on access to the cytosol. Furthermore, we show that PGE_2_ is produced by macrophages and dendritic cells. These data provide new insight as to how CD8^+^ T-cells are primed as well as suggest analysis and modulation of eicosanoid levels, particularly PGE_2_ levels, may be informative to improve the use of *L. monocytogenes*-based immunotherapeutic platforms.

## Materials and Methods

### Ethics statement

This work was carried out in strict accordance with the recommendations in the Guide for the Care and Use of Laboratory Animals of the National Institutes of Health. All protocols were reviewed and approved by the University of Wisconsin-Madison Institutional Animal Care and Use Committee.

### Bacterial strains

The *Listeria monocytogenes* strains used in this study were all in the 10403s background. The attenuated (LADD) strain used in the analysis of T-cell responses was in the *ΔactAΔinlB* background as previously described and engineered to express full length OVA and the B8R_20-27_ epitope[24]. OVA and B8R_20-27_ were constructed as a fusion protein under the control of the *actA* promoter with the secretion signal of the amino terminal 300bp of the ActA gene[13]. This fusion protein was integrated into the site-specific pPL2e vector as previously described[13].

### Mouse strains

Six- to eight-week-old C57BL/6 male and female mice were obtained from the NCI and Charles River NCI facility. *Ptgs2*^-/-^ (COX-2^-/-^) mice were obtained from Jackson Laboratory and maintained as heterozygote breeding pairs. *Ptges^-/-^* (mPGES1^-/-^) mice lacking microsomal PGE synthase have been previously described[67–69]. In order to generate cell-type specific COX-2 knockout mice, COX-2^fl/fl^ mice (stock number 030785) were obtained from Jackson Laboratory and crossed with Lyz2-cre (stock number 004781), CD11c-cre (stock number 008068), or CD4-cre expressing mice (stock number 022071), all also obtained from Jackson Laboratory. Double Lyz2-cre and CD11c-cre expressing mice were generated by crossing COX-2^fl/fl^ Lyz2-cre mice with COX-2^fl/fl^ CD11c-cre mice. Genotypes were confirmed by PCR using the primer pairs in Table 1.

**Table 1.**
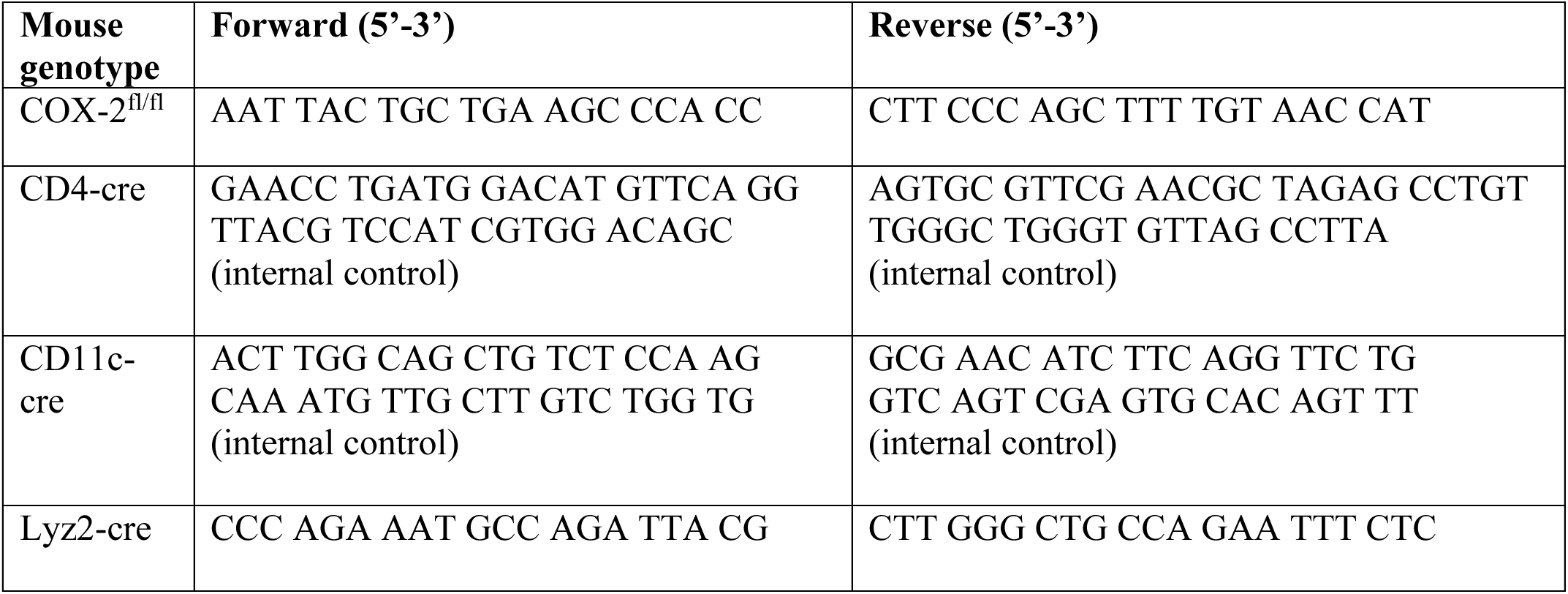
Genotyping primers.

### BMDM and BMDC generation and infection

Bone marrow-derived macrophages and dendritic cells were made using six- to eight-week-old *Ptgs2^-/-^* (COX-2^-/-^) or C57BL/6 mice as previously described[13, 70]. Briefly, bone marrow was harvested and macrophages were cultured in the presence of M-CSF from transfected 3T3 cell supernatant for six days with a supplement of M-CSF at day three and frozen down for storage. Dendritic cells were cultured in the presence of 20ng/ml recombinant GM-CSF (BD Biosciences, San Jose, CA) for 7 days with a supplement of 20ng/mL GM-CSF every third day. For infection, BMDMs or BMDCs were plated at 1×10^6^ cells/well in a 12 well dish overnight +/- 100ng/mL PAM3CSK4. The following morning, cells were infected with indicated strains of *L. monocytogenes* or PBS control at an MOI of 10. Thirty minutes later, supernatant was removed and replaced with medium containing 50µg/mL gentamycin to remove extracellular bacteria. Six hours post infection, cells were harvested for western blot or qRT PCR and supernatant was harvested for eicosanoid analysis as described below.

### qRT PCR

RNA was isolated from BMDMs or BMDCs using the RNAqueous-Micro Total RNA Isolation Kit (Invitrogen), and DNAse treated with Turbo DNAse (Invitrogen) according to manufacturer’s instructions. 500ng total RNA was reverse transcribed in 10µL reactions using the iScript cDNA Synthesis Kit (BioRad) according to manufacturer’s instructions and cDNA was diluted 10-fold using molecular grade water (Invitrogen). 2.5µL diluted cDNA was used as template in a 10µL qRT-PCR reaction performed in duplicate using gene-specific primers and Kapa SYBR Green Universal qPCR mix (KAPA Biosystems) according to manufacturer’s instructions using a BioRad CFX Connect Real-Time PCR System. The sequences of gene-specific primers are shown in Table 2. Data was analyzed using Excel and all RNA abundances were calculated by using a standard curve of synthesized template (Integrated DNA Technologies, G-Blocks) and are normalized to *ActB* (β-actin).

**Table 2.**
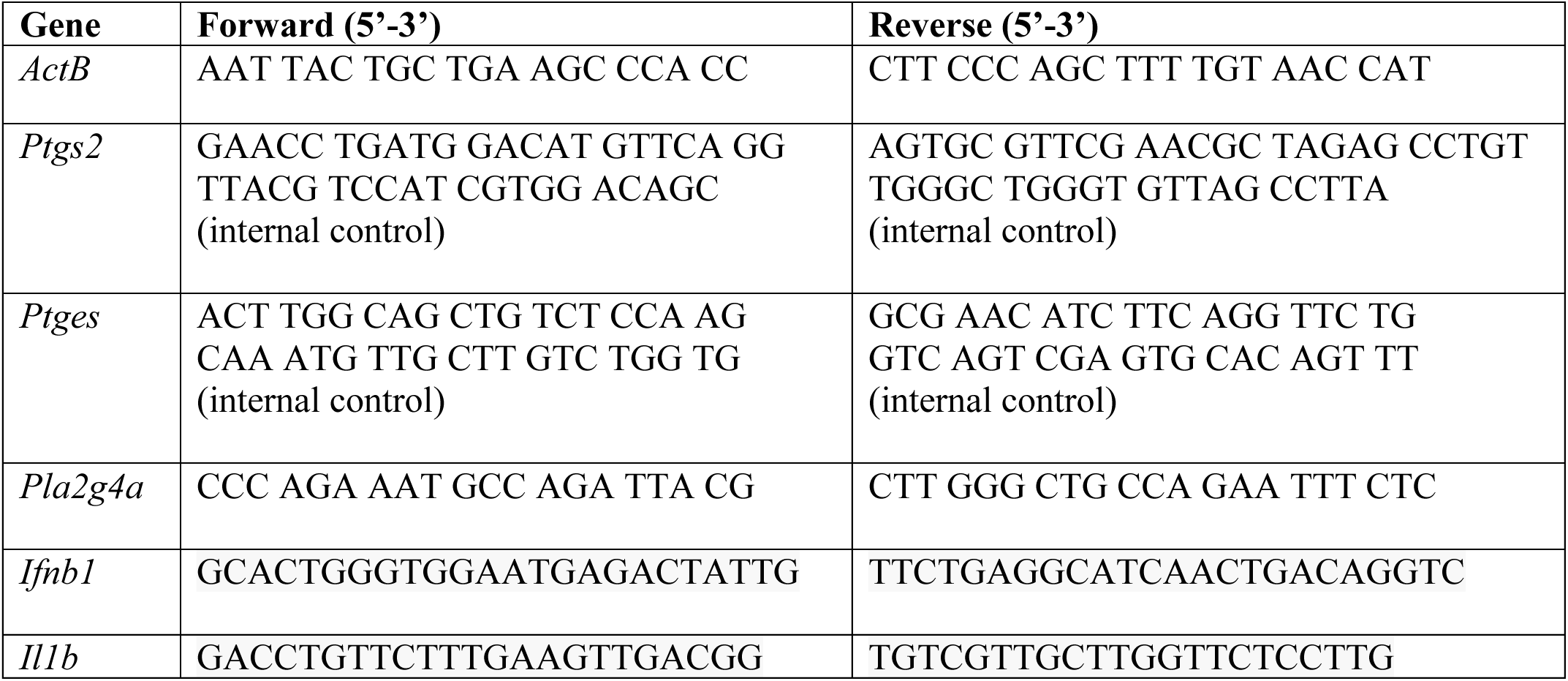
qRT PCR primers.

### Western blots

BMDMs or BMDCs were harvested and protein was extracted using the Pierce SDS-PAGE Sample Prep Kit (Thermo) according to the manufacturer’s instructions. Total protein content was measured by the Pierce BCA Protein Assay Kit (Thermo) and equivalent protein levels were loaded into an polyacrylamide gel (BioRad). Samples were transferred onto a nitrocellulose membrane using a semi-dry transfer apparatus before blocking with a 5% skim milk solution for thirty minutes at room temperature. After washing 3x with PBS-T, the membrane was incubated overnight at 4°C with the primary antibodies anti-COX-2 (1:200, Cayman Chemical) and anti-β-actin loading control (1:1000, ThermoFisher) in a 5% bovine serum albumin solution. The following day samples were washed with PBS-T before being incubated with secondary antibodies (anti-rabbit 800 at 1:10,000, anti-mouse 680 at 1:5,000). Samples were imaged on a LiCor imager and analyzed via ImageStudio. Sample signal was normalized to β-actin and relative abundance was compared to wild-type *L. monocytogenes*.

### *In vivo* immunizations and pharmacological treatments

*L. monocytogenes* of the wild-type, attenuated (LADD), or *Δhly* background were grown overnight in brain heart infusion media at 30C. The bacteria were back diluted 1:5 and allowed to grow to log phase (OD0.4-0.6, ∼1-1.5 hours) at 37C, with aeration, prior to infection. Bacteria were diluted in PBS and mice were infected with 200µL at the indicated doses intravenously. For bacterial burden analysis, mice were sacrificed at 12hpi and livers were homogenized in 0.1% Nonidet P-40 in PBS and plated on Luria-Bertani plates. For splenic macrophage depletion, 200µL clodronate, PBS control, or endosome lipid control (Encapsula Nano Sciences) were given intravenously 24 hours prior to bacterial infection according to the manufacturer’s instructions. Depletion efficacy was confirmed by assessing abundance of splenic CD11b^+^ cells (clone M1/70) and CD11c^+^ cells (clone N418) by flow cytometry. Celecoxib (Cayman Chemical) was milled into standard mouse chow (Envigo) at 100mg/kg and fed ad lib for 48 hours before and after immunization[14, 71].

### Eicosanoid measurement

For *in vivo* eicosanoid extractions, spleens from mice were harvested at twelve hours post immunization and flash frozen in tubes containing 50ng deuterated PGE_2_ standard (Cayman Chemical) in 5μL methanol and stored overnight at -80C. For *ex vivo* extractions, 1mL of supernatant was flash frozen in tubes similarly containing 50ng deuterated PGE_2_ standard in 5μL methanol. The following day, two mL of ice cold methanol were added to the tissue culture supernatant or spleens. Spleens were homogenized in glass homogenizers. Samples then were incubated at 4C for 30 minutes. Next, cellular debris was removed by centrifugation and samples were concentrated to 1mL volume before being acidified with pH 3.5 water and loaded onto conditioned solid phase C18 cartridges. Samples were washed with hexanes before eluting using methyl formate followed by methanol. Samples were concentrated using a steady stream of nitrogen gas and suspended into 55:45:0.1 MeOH:H_2_O:acetic acid and analyzed on an HPLC coupled to a mass spectrometer (Q Exactive; Thermo Scientific) using a C18 Acquity BEH column (100mm x 2.1 mm x 1.7μm) operated in negative ionization mode. Samples were eluted with a mobile phase 55:45:0.1 MeOH:H_2_O:acetic acid shifted to 98:2:0.1 over 20 minutes. Mass-to-charge ratios included were between 100 and 800 and compared to standards (Cayman Chemical) by analysis via MAVEN.

### T-cell analysis

Mice were sacrificed seven days after immunization and splenocytes were isolated as previously described[4]. In brief, red blood cells were lysed using ACK buffer and then splenocytes were counted using a Z1 Coulter counter. For tetramer analysis, splenocytes were immediately blocked for Fc (Tonbo Bioscience) and stained for B8R tetramer (AF488, 1:300, NIH Tetramer Facility, Atlanta, GA) followed by staining with anti-CD3 (PeCy7, 1:100, clone 145-2C11) and anti-CD8*α* (eFlour450, 1:200, clone 145-2C11). Cells were then stored overnight at 4C in a 1:1 of IC Fixation Buffer (ThermoFisher Scientific) and FACS buffer. For analysis of cytokine production, 1.7×10^6^ cells were plated in a 96 well dish and incubated for five hours in the presence of B8R_20-27_ (TSYKFESV) or OVA_257-264_ (SIINFEKL) peptides and brefeldin A (eBioscience). Splenocytes were then subjected to FC block (Tonbo Bioscience) and stained with anti-CD3 (FITC, 1:200, clone 145-2C11) and anti-CD8*α* (eFlour450, 1:200, clone 53-6.7) before treatment with fixing and permeabilization buffers (eBioscience). Cells were then further stained with anti-IFN*γ* (APC, 1:300, clone XMG1.2). Samples were acquired using the LSRII flow cytometer (BD Biosciences, San Jose, CA) and analyzed with FlowJo software (Tree Star, Ashland, OR).

### Cryosection preparation and immunofluorescence microscopy of infected spleens

C57BL/6 mice were infected intravenously by tail vein injection of 10^7^ LADD *L. monocytogenes* in 150 μl of PBS. Mice were sacrificed at 3 or 10 hpi and spleens harvested and snap frozen in OCT for immunofluorescence microscopy as described previously[20]. Uninfected mice were used as negative controls. Briefly, 5μm spleen cryosections were cut using a Leica CM1850 cryostat, mounted on Superfrost Plus microscope slides (Thermo Fisher) and stored at -80°C until use. Slides were fixed in 10% buffered formalin phosphate at RT for 5 minutes and sections, washed in TBS and blocked with StartingBlock T20 Blocking buffer containing Fc blocker (Thermo Scientific, 37543). Sections were incubated with unconjugated *L. monocytogenes* monoclonal Ab (Invitrogen, MA1-20271), anti-MARCO polyclonal Goat IgG-Biotin (R&D Systems, BAF2956), and FITC-conjugated COX2 polyclonal antibody (Cayman Chemical, 10010096) at 1/100-200 dilution at RT for 1-2h in dark humidified incubation chamber or isotype control antibodies including Rabbit IgG-FITC (Invitrogen, 11-4614-80), Armenian Hamter IgG-PE (Invitrogen, 13-4888-81) and Mouse IgG2a kappa (Invitrogen, 14-4724-81). Biotinylated and unconjugated primary antibodies were detected by incubating with Streptavidin-PE (Pharmingen, 534061) and Rat anti-mouse IgG2a antibody (Invitrogen, 17-4210-80) respectively. Slides were preserved using ProLong Diamond Antifade mounting media (Invitrogen, P36965) and clear nail polish to seal the edges. Slides were analyzed using an Olympus IX51 fluorescence microscope equipped with LCPlanFL 20x/0.4 NA and UPlanFL 40x/1.3 oil objectives, an X-Cite 120 excitation unit (Exfo), FITC/PE/APC optimized filter sets (Semrock), an Orca Flash 2.8 monochrome camera (Hamamatsu) and SlideBook software (Intelligent Imaging Innovations) for hardware control and image acquisition. Images were captured with both 20x an 40x objective with exposure times ranging from 200-400ms. Pseudo-colored 3-channel RGB mages were imported into Imaris 9.6 (Bitplane) for smoothing, contrast enhancement (linear contrast stretch), annotation and colocalization analysis.

### Statistical analysis

Statistical analysis was performed by GraphPad Prism Software (La Jolla, CA) and analyzed via Mann Whitney *U* test or one-way ANOVA with Bonferroni’s correction as indicated.

## Acknowledgments

We would like to thank the NIH Tetramer Core Facility for provision of MHC-I B8R tetramers.

